# Predicting Transcriptional Regulatory Activities with Deep Convolutional Networks

**DOI:** 10.1101/099879

**Authors:** Joe Paggi, Andrew Lamb, Kevin Tian, Irving Hsu, Pierre-Louis Cedoz, Prasad Kawthekar

## Abstract

Massively parallel reporter assays (MPRAs) are a method to probe the effects of short sequences on transcriptional regulation activity. In a MPRA, short sequences are extracted from suspected regulatory regions, inserted into reporter plasmids, transfected into cell-types of interest, and the transcriptional activity of each reporter is assayed. Recently, Ernst et al. presented MPRA data covering 15750 putative regulatory regions. We trained a multitask convolutional neural network architecture using these sequence expression readouts which predicts as output the expression level outputs across four combinations of cell types and promoters. The model allows for the assigning of importance scores to each base through *in silico* mutagenesis, and the resulting importance scores correlated well with regions enriched for conservation and transcription factor binding.

## 1 Introduction

The assignment of significance values to bases across different positions along the genome is a problem which has many applications in studies of various genetic functions, including genome-wide association studies (GWAS), with respect to gene regulation and disease for example. The general problems that we aim to consider are the following: (1) for a particular sequence or region, how well does that sequence or region act as an enhancer for downstream gene expression, and (2) for a given position along the genome, how can we assign some “importance score” as to how significant that particular base is in determining gene expression and other related functional cell processes? A method which is able to successfully yield insight towards these questions can be used in discovering bases or SNPs which affect cell function, for example disease-causing SNPs.

The most direct means to assess the importance of sequences in cell function regulation is to measure its success in causing expression in a reporter assay. Recently, Ernst et al. in their Sharpr-MPRA study released a large amount of data which were the results of performing multiple reporter assay experiments on over 4 million nucleotides coming from various putative regulatory regions [3]. The main novelty factor of these experiments was in their scale and their resolution - the sequences assayed were tiled 5 base pairs (bp) apart, so the relative contributions of positions could be determined at a higher resolution. The authors developed a method using a probabilistic graphical model to assign contributions to positions.

We believe that the richness in the MPRA dataset can be better exploited using a deep learning model which takes as input a sequence and produces scores predicting reporter assay expression reads in the same experiments as were performed in the MPRA scale-up study. For one thing, if done successfully this would allow for imputed expression predictions on sequences which were not in the original study. Furthermore, this would allow for more natural importance scoring methods for bases, because their predicted contribution to the expression level can be tested by perturbation, for example. It also extends these importance scoring methods to bases not in the training set.

## 2 Related Work

### 2.1 Ernst et al. Computational Model

#### 2.1.1 Description of Model

Ernst et. al. developed a probabilistic graphical model (PGM) to predict the regulatory activity of each base from their assay. First, the model maps variables corresponding to the regulatory activity of each of 59 5-bp intervals (the tile length) to the 31 reporter assay measurements for each 295-mer. Then, their method modeled the importance of each individual base by linear interpretation, which is a relatively restricting assumption. Indeed, without some prior assumptions on the relationship between relative contributions of bases in each 5-bp tile, the PGM cannot assign basewise importance scores after computing scores for each tile variable.

The goal of the Sharpr-MPRA model was to infer the activity of each nucleotide by using the activity of the tiles as a proxy, and they made no attempt to make a model mapping from sequence to activity, which is fundamentally distinct from our proposed approach. For these reasons we feel their model can be improved upon.

#### 2.1.2 Results

The results on the regulatory effects of nucleotides were validated by Ernst et. al. through comparison with various existing results. Agreement with CENTIPEDE motif annotations and evolutionarily conserved elements supported the success of Sharpr-MRNA in discovering functional nucleotides. The method agreed with DeltaSVM predictions on activating nucleotides, but also discovered repressive nucleotides which the former did not. The success of higher-density tilings also was demonstrated through sparsity experiments. Among interesting new ideas captured were the fact that the most highly active nucleotides were enriched in retroviral repeat elements, and that across cell types the strongest activity sequence pattern was a motif not associated with an established regulator.

## 3 Dataset and Pre-Processing

The input of our model will be a sequence of 145 bp for every region and every tile. There are 15,720 regions and 31 tiles per region, and 2 replicates, for a total of approximately 980,000 examples.

The label will be the normalized activation scores calculated from the DNA count and the mRNA count, from SHARPR. In the original study, each sequence yielded 8 activation scores since it was measured twice for 2 different cell types and for 2 promoters. We consider an experiment to be a (cell type, promoter) pair, and define the error of our model to be the mean squared error of our predicted expression levels with respect to the average expression level across two replicates for each (experiment, sequence) pair. The models we considered for predicting these expression scores were multi-task neural nets, which simultaneously produced these 4 predicted experiment activation scores from an input sequence.

One difficulty that this method inherently has is the amount of experimental noise present in the dataset. The amount of correlation between the two replicates was often surprisingly low, but was much more noticeable when neither read was very high. For the four experiment combinations of a cell type and a promoter, the correlations between replicates ranged from 0.325427 to 0.467690.

We intuited that the reason for this was that for sequences which were strongly enhancing or repressing, the reads would be more consistent because they are not as susceptible to experimental noise, but for other sequences the reads would be more inconsistent. This was the motivation for weighting our error function with respect to the consistency between reads, as discussed in Section 4.2.3.

### 3.1 Pilot Study

A pilot study was designed to test the tiling approach for reproducibility and cell-type specificity of predicting regulatory function of nucleotides, and to compare the accuracy of predictions with known endogenous epigenomic information. 250 regulatory regions were tested across two cell types (K562, HepG2) using a SV40 promoter. Each regulatory region consisted of 385 nucleotide regions that were tiled with 145 nucleotide constructs, each with 24 unique barcodes, at 30 nucleotide steps. The observed level and frequency of normalized regulatory (MPRA) activity for the sequences were observed to correspond with endogenous epigenomic information, were reliable across replicates, and were cell-specific as the nucleotide differences between neighboring tiles were found to correspond to cell-specific motifs.

### 3.2 Large-scale Study

For the large scale experiments, the most informative 295-bp positions from the 385-bp regions tested in the pilot study were obtained from 15,720 putative regulatory regions retrieved from 4 different cell types (H1-hESC, HepG2, K562, HUVEC). These regions were tiled with 145-bp nucleotides at steps of 5-bp, with a single barcode attached to each construct. Each tile was replicated and tested in two of the cell types (HepG2, K562) with two promoters each (minP, SV40P) across two spot arrays, for a total of 16 experiments. The DNA and RNA counts for each assay were smoothed (by adding 1 to the counts) and normalized (by dividing each smoothed count by the sum of all smoothed counts), and the log base two ratio for the normalized smoothed counts was calculated. This log-ratio was held as a measure for the direction (+/-; activation/repression) and magnitude of regulatory function for the region tested. The sequences that didn’t have accurate values for all experiments as determined by the SHARPR software were filtered out. For the purposes of developing our model, we trained only on the experiment reads from this scale-up study, but as previously mentioned having less replicates of each experiment made these expression values more susceptible to noise.

## 4 Methodology

### 4.1 Baseline *k*-mer Model

In order to provide a baseline to which to compare our CNN, we developed a *k*-mer based linear regression model. The model takes counts of the number of occurrences of all possible 6-nucleotide sequences as input features and from these predicts the output of the MPRA. The linear regression model used was the stochastic gradient decent linear regression model from sklearn. We chose *k* = 6 because it outperformed smaller *k* (data not shown) and we were unable to run larger values of *k* because of memory restrictions.

### 4.2 Multi-Task Neural Nets

Our goal is to obtain reporter activity of nucleotide regions at single-nucleotide resolution. The data from the original study maps 145-bp sequences to MPRA activity scores for different cell type and promoter combinations. In order to obtain nucleotide-level resolution of activity scores, we first train a regression model to accurately learn this mapping for each task from the experimental data, and then use state-of-the-art feature importance scoring techniques to identify the effect of each nucleotide for all 145-bp sequences.

We use multi-task convolutional neural networks as our predictive model since we expect the activity scores of a 145-bp sequence to be correlated across across each cell-promoter combination. Multi-task ConvNets share parameters across some layers, which then lead up to task-specific layers. This parameter sharing in the base network can be useful to counter overfitting by forcing generalization across all tasks.

#### 4.2.1 Input and Output

The inputs to our model are one-hot-encoded 145-bp nucleotide sequences. The corresponding outputs for each input are the average activity scores across the two replicate measurements for each of the 4 tasks (cell-type and promoter-type combination).

We make a minor modification to the one-hot-encoded input sequences based on the experimental setup of the original study. Specifically, the authors of the original study sampled the 295-bp nucleotide regions such that they are likely to be centered at known motifs. This is shown in Figure 1 obtained from 2.1, which plots the average absolute activity scores by position across all 295-bp regions. Consequently, since the position of nucleotides are correlated with their relative activity, we pad each one-hot encoded vector of 145-bp sequences with 0s on either side of the tile’s position in the region to obtain a (295,4)-dimensional one-hot encoded vector for each tile in the region (Figure 2).

**Figure 1:**
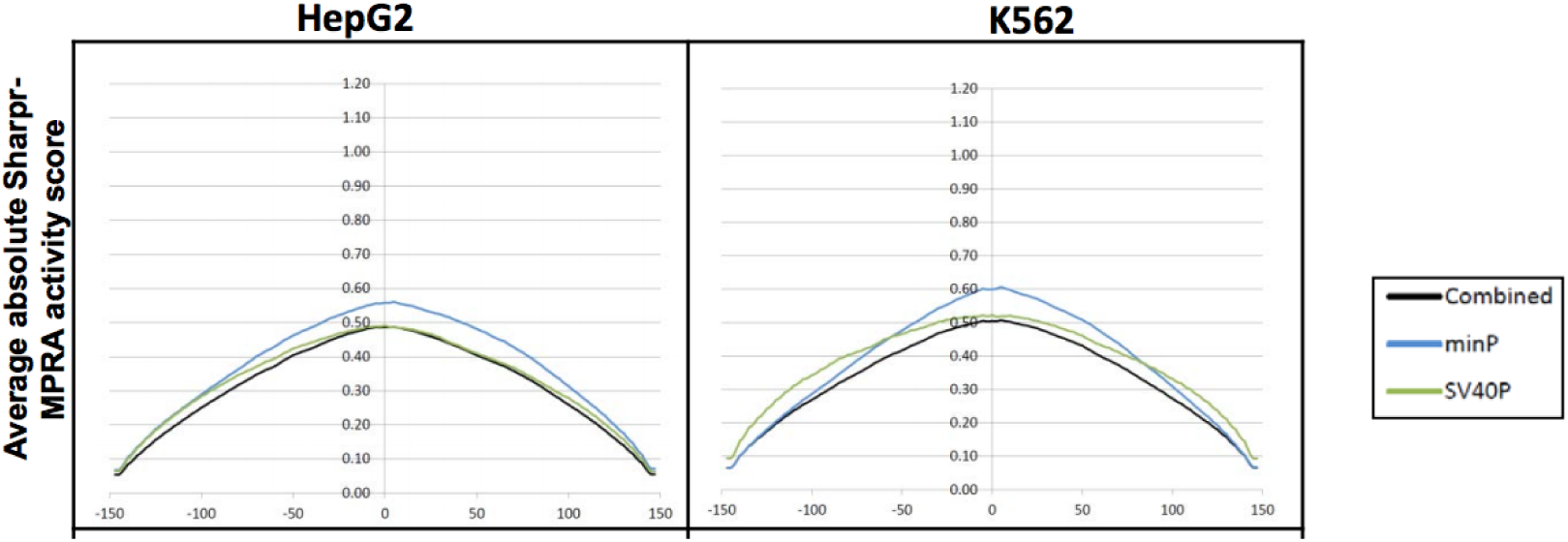
Sharpr-MPRA average absolute activity Scores by position.

**Figure 2:**
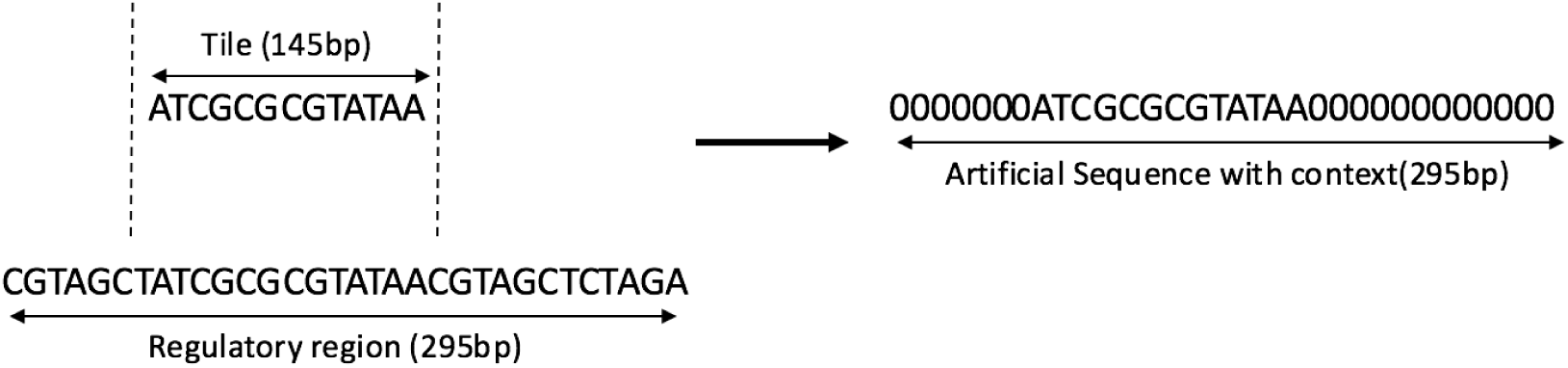
Artificial sequence to account for the position of the tile in the regulatory region.

#### 4.2.2 Training and Fine-Tuning

We implement our model in DragoNN [1] and train it to minimize the average MSE of prediction across all tasks. We use the default multi-task net architecture of DragoNN with hyperparameter tuning along the following variation points-

- Number of convolutional Layers: 1-10
- Number of dense layers: 1-2
- Number of filter per conv layer: 30-100
- Conv width: 8-15
- Pool width: 10-50
- Dropout: 0.1

The best performing model consists of 2 convolutional layers each with ReLU activation, 100 filters, conv-width of 4, and dropout probability of 0.1. The convolutional layers are followed by a max-pooling layer with pool-width of 44 and a dense layer also with ReLU activation and 4 outputs corresponding to the four tasks. The weights of the layers are initialized to Gaussian scaled by fan-in [4]. We also validate different optimizers and select Adam as it shows signs of faster convergence [2]. The final architecture is shown in Figure 3.

**Figure 3:**
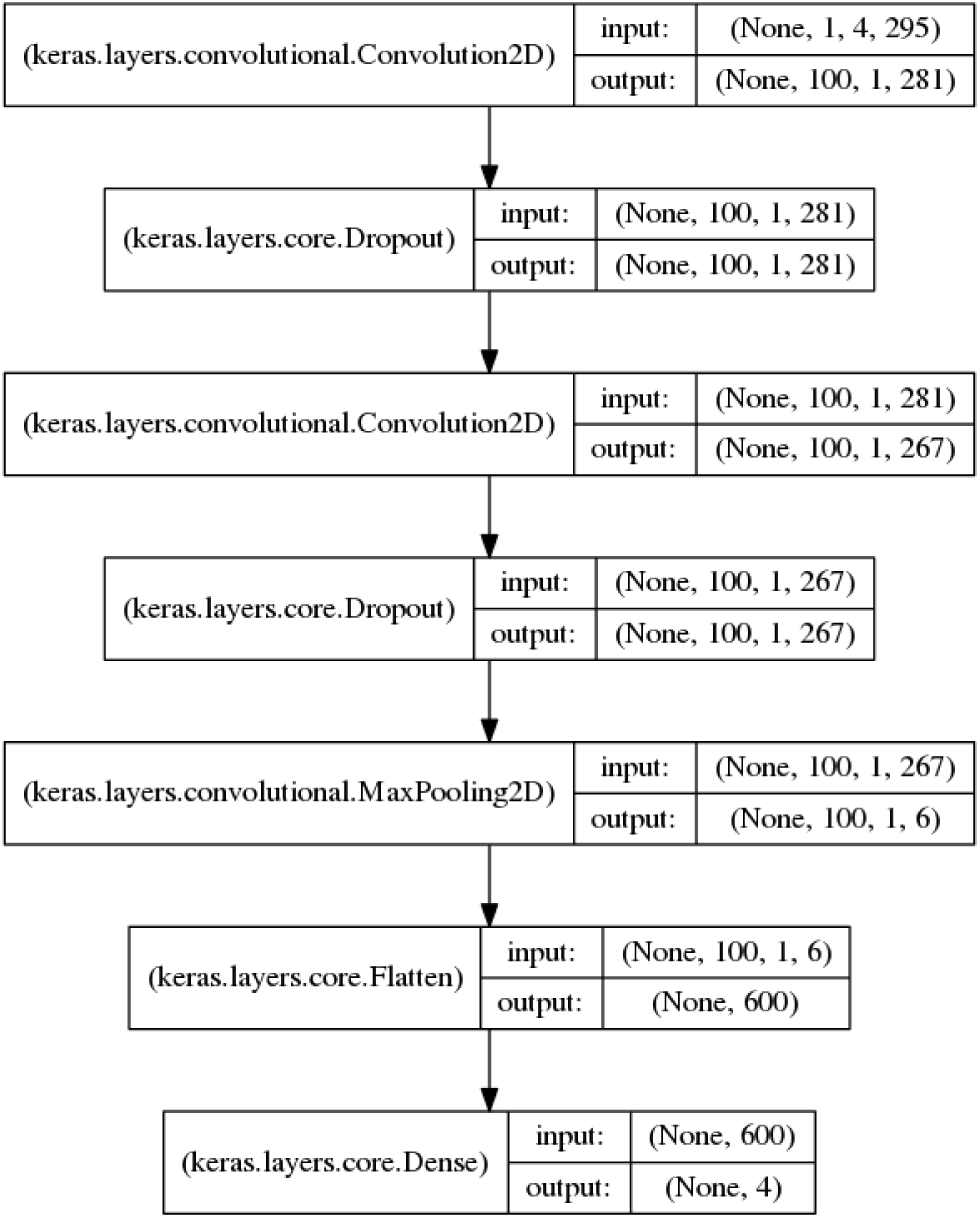
295 bp sequences input.

**Figure 4:**
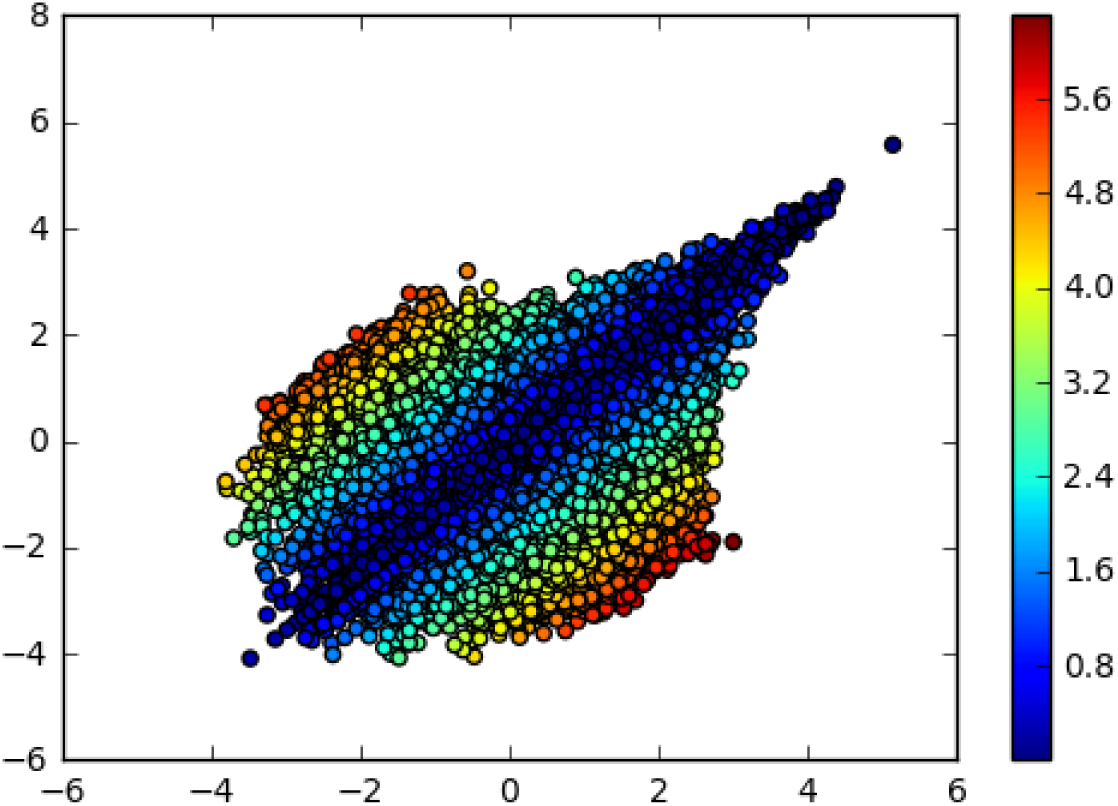
Distance between replicates for (HepG2, MinP) task.

##### Weighted Inputs

To account for variance in the experimental measurements, we weight the input sequences during training based on the dissimilarity of the MPRA measurements across the two replicates. The weight of an example is the negative distance of the replicates scaled between 0 and 1, such that reproducible measurements receive larger weights.

The MSE learning curves for the best-performing architecture on all tasks (and on the mean for all tasks) is shown in 5

**Figure 5:**
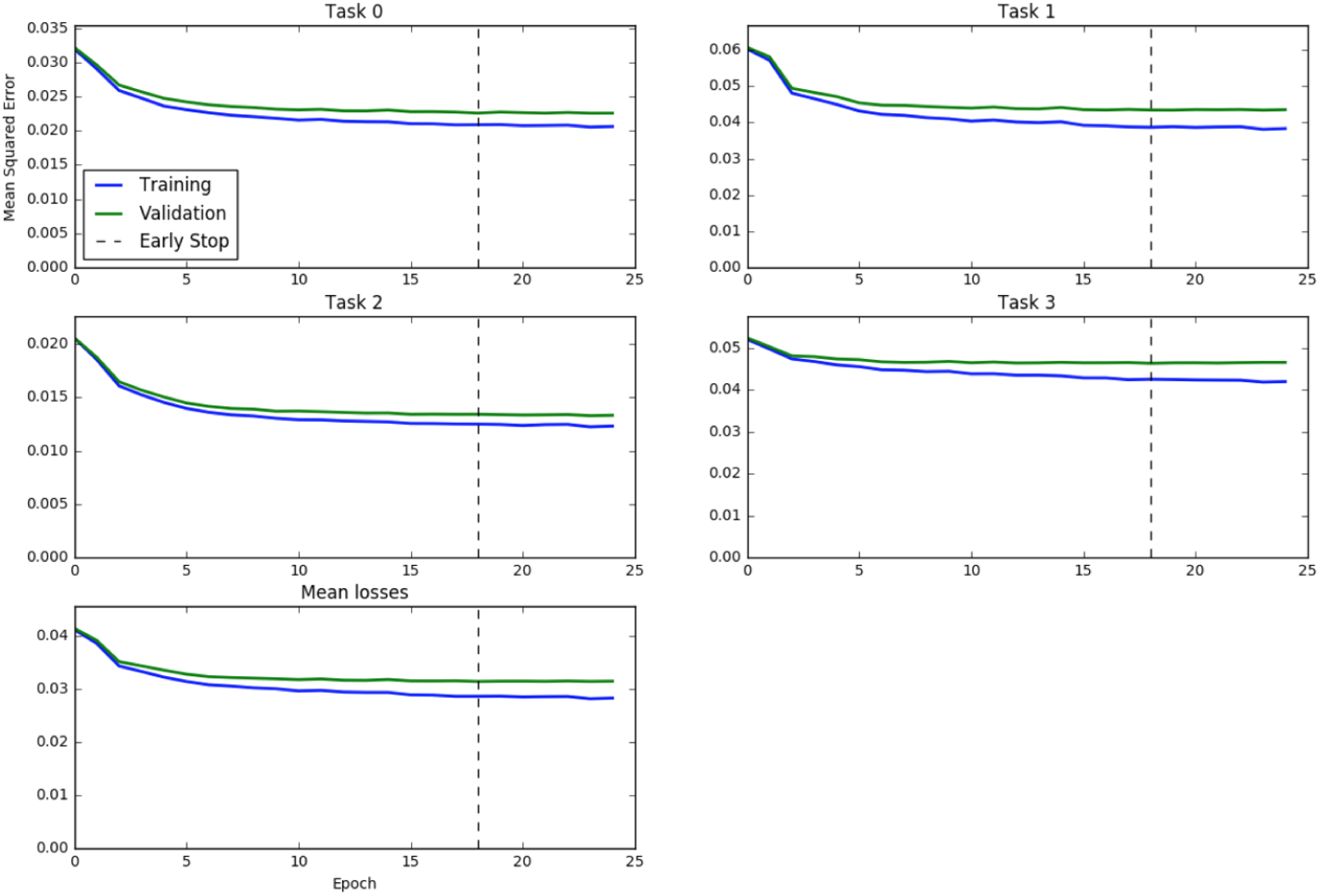
MSE Learning curves for best-performing architecture on all tasks and on mean for all tasks. Task 0 corresponds to (HepG2, minP) combination, task 1 to (K562, minP), task 2 to (HepG2, SV40P), and task 3 to (K562, SV40P).

**Figure 6:**
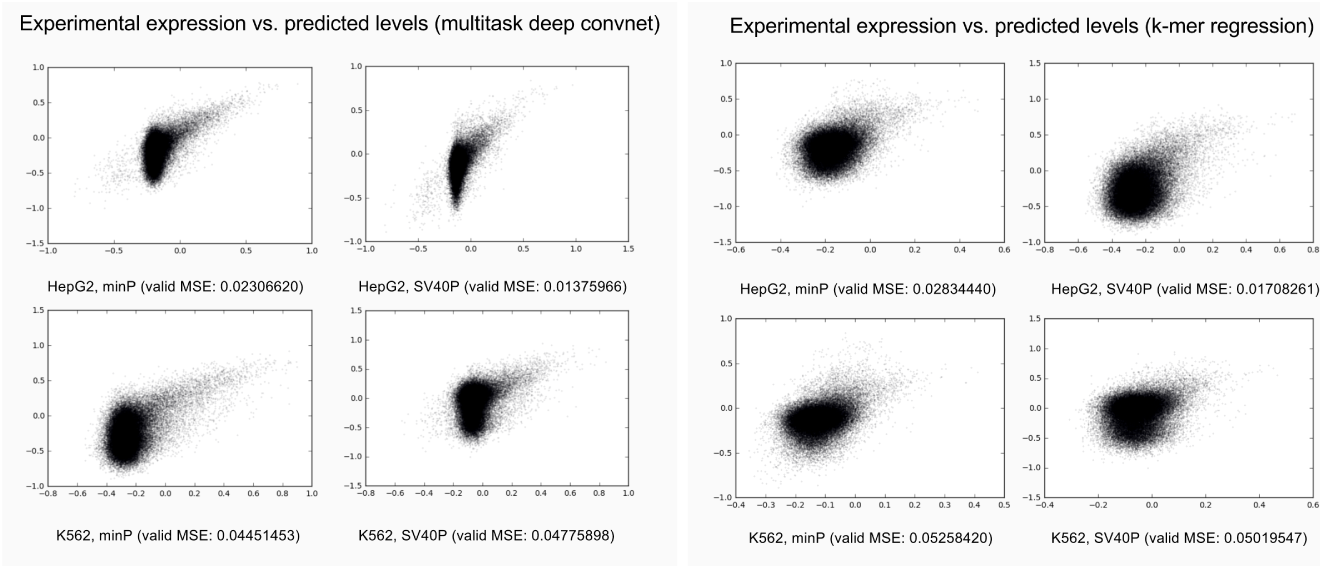
Our multitask CNN outperforms k-mer based linear regression.

### 4.3 Interpretation of the Trained Network

A major goal of our work was prediction of activation and repression function at a single nucleotide level. During training, the parameters of the model are tuned to predict the reporter activity for the 145-bp sequences in the the training set, so additional computational techniques are needed to obtain single nucleotide resolution. Sharpr-MPRA used a probabilistic graphical model, where the activity for 5-bp intervals were modelled as hidden variables, and regulatory activity for single nucleotides was inferred with linear interpolation [3]. Our model allows us to slide the sequence which we perform reporter assay expression predictions (and subsequently compute importance scores), so that these scores can be calculated in the context of sequences which weren’t perfectly aligned with the 145-bp sequences for which experimental results were obtained, something not allowed by the Sharpr-MPRA model. We took advantage of this fact in our importance computations.

#### 4.3.1 *In Silico* Mutagenesis

In *in silico* mutagenesis (ISM), a form of perturbation testing, each individual base location is perturbed (changed into every other possible base) and the corresponding predicted expression level readout is changed as a result. This produces 4 scores (at most 3 of which are nonzero), computed as how much more expression results from the sequence with original base than with the perturbed base. We scored the importance of the base as the (signed) difference with maximum absolute value. The intuition is that for bases important to enhancing expression, this value will be a large positive, and for bases important to repressing it will be a large negative (it will otherwise be relatively small).

The method is relatively computationally efficient, as for each sequence it requires a number of forward passes linear in the number of the length of the input sequence and yields an importance score for each base. The (unsigned) importances should correlate positively with those assigned by SHARPR for positions where scores are available. We also expected base positions with large ISM importance values to be enriched in conserved regions and regions with known transcription factor binding.

We experimented with a few different methods in order to normalize well, because the distribution of bases in the dataset was uneven. In particular, bases which lie closer to the edges of a 295-bp region will have less predicted scores associated with it, because it belongs to less 145-mers which were assayed. We circumvented this problem by taking advantage of the flexibility of our model to extend to sequences which were not assayed.

The scoring method we used consisted of computing 5 importance scores for each nucleotide position in a 295-bp sequence. We did so by passing 15 145-mers which were tiled every 29 bp into our model, such that each position in the original 295-mer was in exactly 5 of these 145-mers. For example, if the indices of the original 295-mer are 0 through 294, the indices of the sequence we performed predicted assays for were (−116 through 28), (−87 through 57), … (290 through 434). Thus, each base will have one predicted importance score in each quantile of the sequence, which should correct for any inherent bias the importance scoring has with respect to location in the input sequence. We in fact observed better correlation under the metrics discussed below with this method, which led us to believe it was a more accurate way for determining the scores for each genomic position.

### 4.4 Alternative Importance Scoring Methods

Among the other methods we tried in order to interpret our model was the DeepLIFT method proposed by Shrikumar et. al. [7] in assigning importance scores to the input features through propagating activation differences. However, we were computationally constrained and ended up choosing ISM as our method for determining these scores. In the future, we would like to experiment more with alternative scoring methods, including gradient-based methods, to see if they show higher levels of agreement with our evaluation metrics and serve as a better way to perform this score assignment.

## 5 Results

### 5.1 Multitask CNN Outperforms K-mer Regression

We evaluated the performance of our model by comparing its performance to a k-mer based linear regression method (methods). We found that our CNN performs 10 to 15 % better for all tasks. In addition to outperforming the baseline model, it is apparent from a plot of predicted score to experimental assay output that our model captured much of the meaningful variation in the data.

### 5.2 Importance Scores Correlate with SHARPR Scores, Conservation, and TF Binding Sites

In order to prove the efficacy of our nucleotide resolution activity scores, we examined the correlation between these scores with SHARPR scores, sequence conservation, and known TF binding sites. We found that our scores were significantly correlated with the SHARPR scores 7.

**Figure 7:**
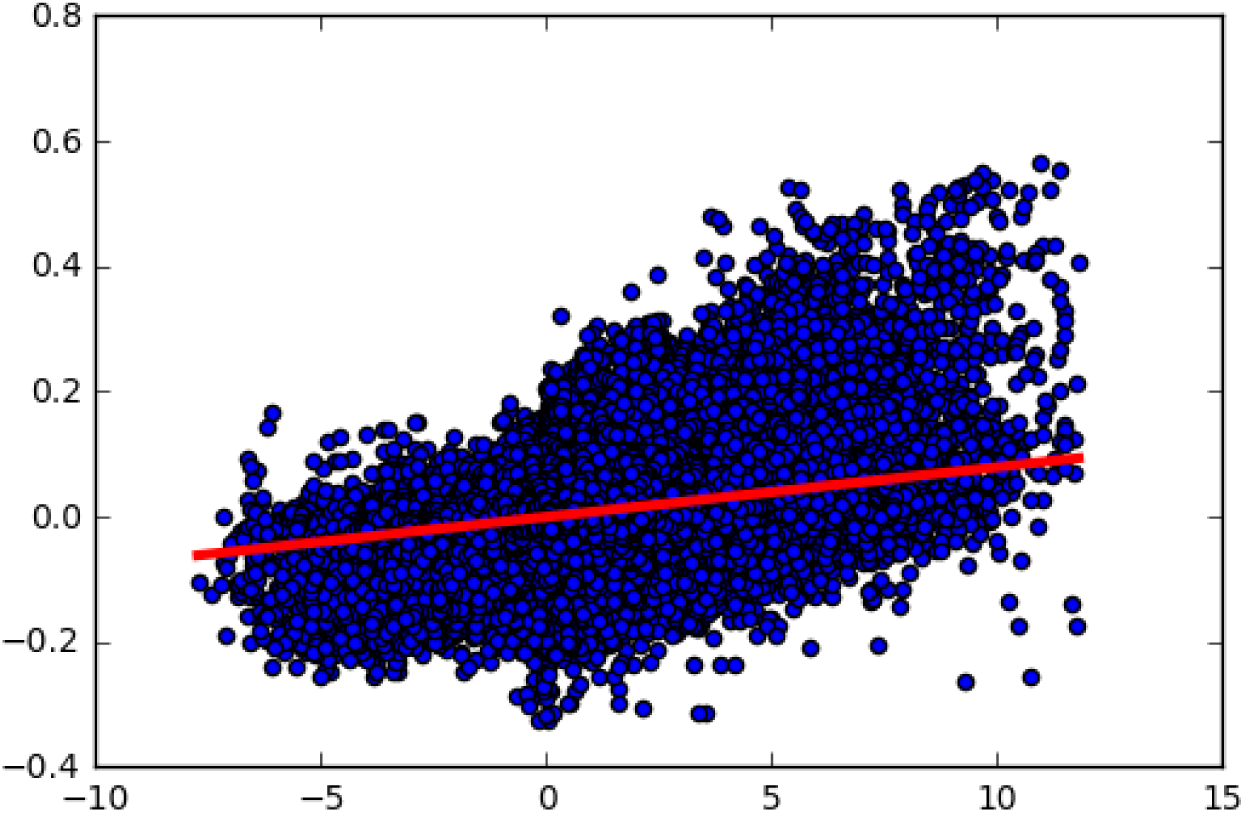
SHARPR scores (x-axis) are correlated with a Spearman Rank Coefficient of 0.16 to ISM scores (y-axis) (statistically significant with *p* ≈ 0).

We assessed the correlation between our scores and SiPhy, a dataset of conserved regions calculated from the genomes of 29 mammals [5]. We divided nucleotides into 5000 quantiles of activity scores. For each quantile, we calculated the fraction of nucleotides that were reported as conserved. We found that quantiles with both a positive and negative activity score tended to have an increased overlap with conserved nucleotides 8.

**Figure 8:**
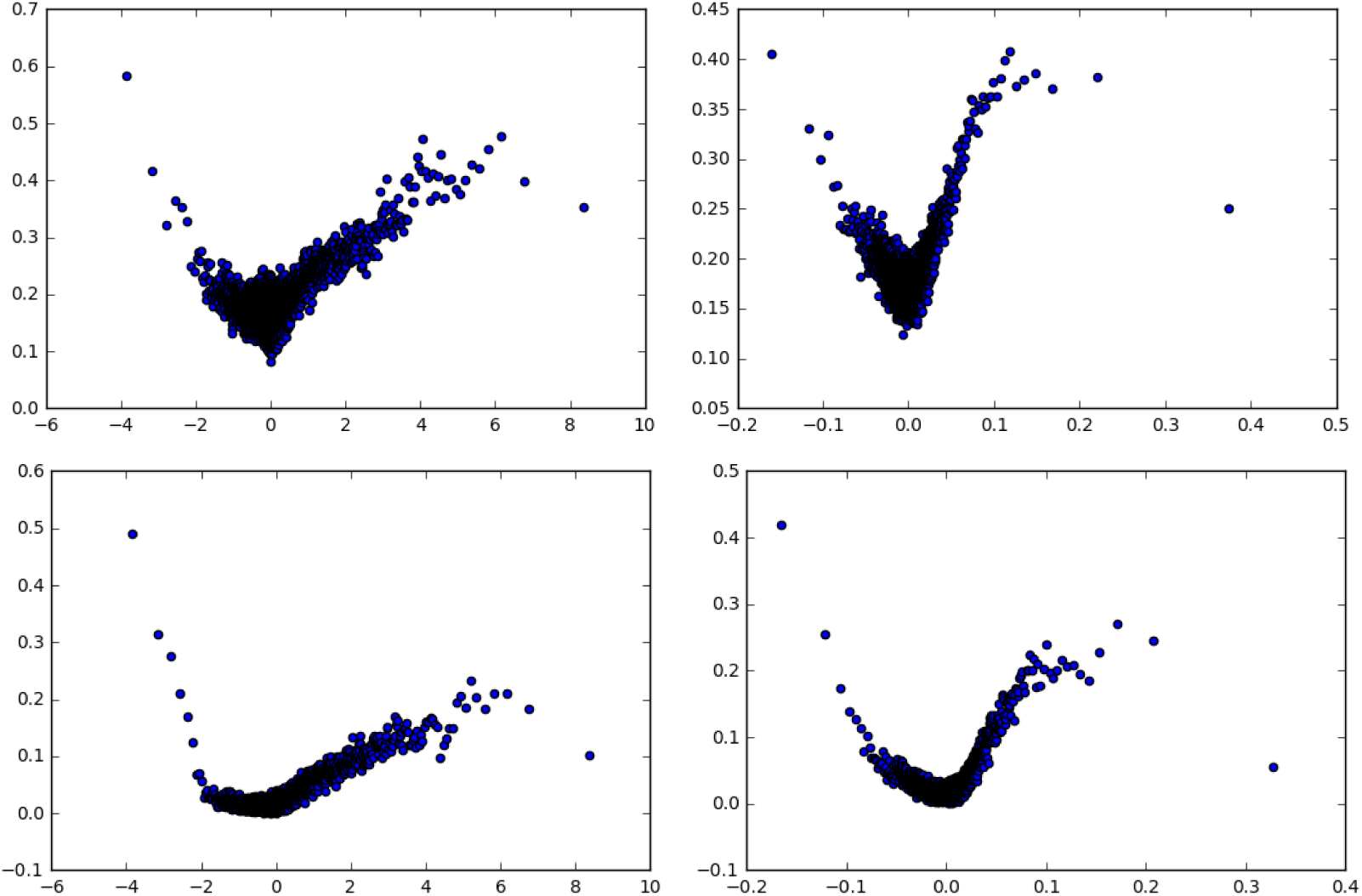
ISM scores (right) show a strong correlation with SiPhy conservation scores (top) and Centipede TF binding sites (bottom). Equivalent SHARPR scores are shown on the left. All data is for HepG2 cells with a minimal promoter.

We performed a similar analysis for overlap with CENTIPEDE cell-type-specific TF binding sites [6]. We again found that quantiles with both positive and negative activity scores show an increased overlap with TF binding sites 8.

### 5.3 Example Applications

To show the utility of our method for discovering biologically meaningful trends, here we present two example applications. First, we show a plot of the average regulatory score for all transcription factor binding sites predicted by Centipede. We see that the scores correlate well between cell types, but that there are some that show significantly more activity in one cell type than the other. Next, we asked a simple question: Does enhancer density correlate with distance from promoters? To do this we prepared a meta-plot of regulatory activity versus distance upstream of a promoter. We found that directly upstream of promoters is highly enriched for enhancers and repressors are excluded.

**Figure 9:**
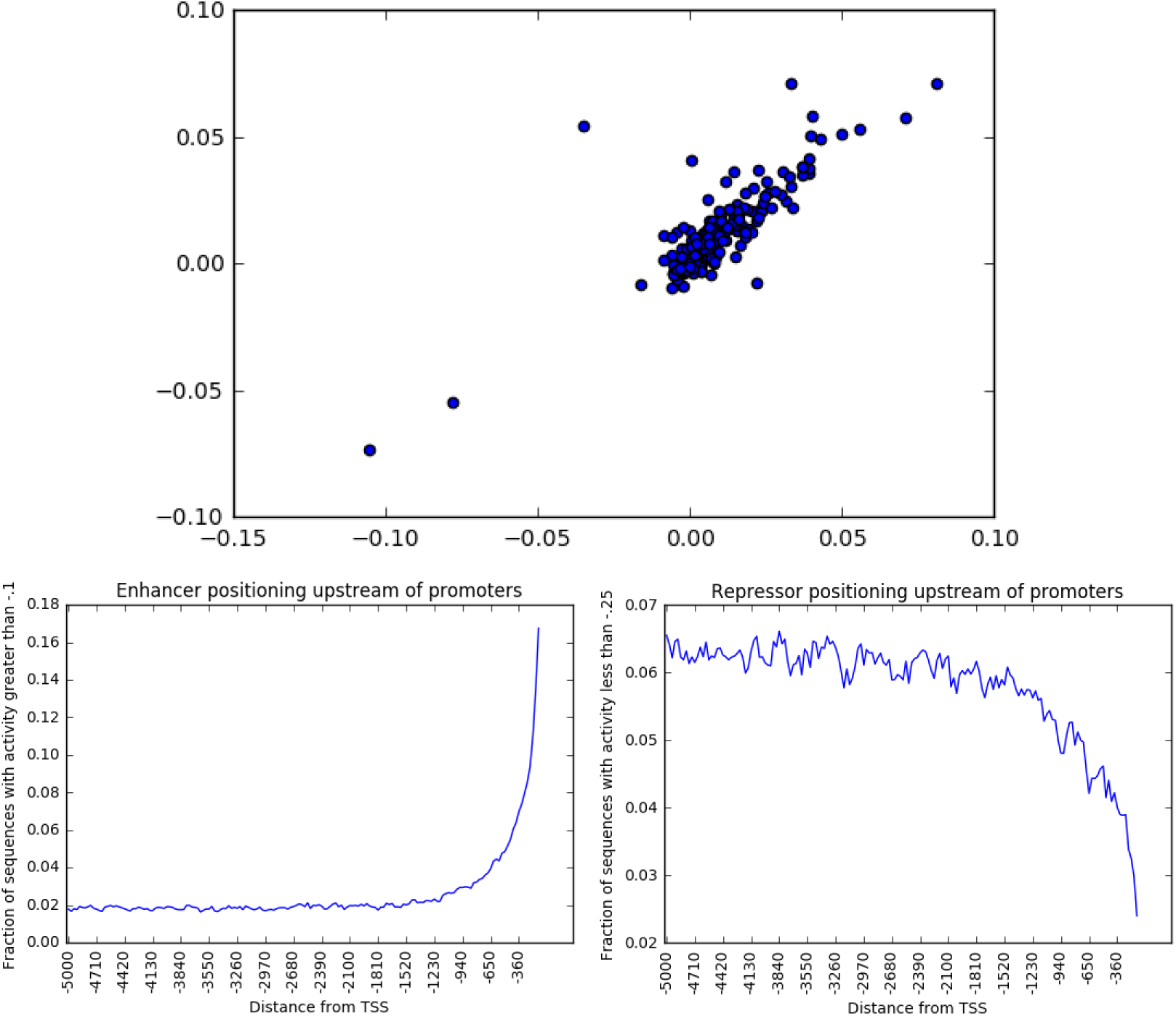
Example applications of our model. On top is average regulatory score for TFs in Centipede dataset in HepG2 versus k562. Bottom shows density of enhancers and repressors as a function of distance upstream of a promoter.

## 6 Future Work

There are a few additional directions in which we wish to take this study. Perhaps foremost is experimenting with additional ways to generate importance scores given our model, to see if they are more predictive or natural, as mentioned previously.

We would like to see how well our predictions or the general idea of this family of deep models generalizes to other locations along the genome, which did not belong to the regions directly assayed by the Sharpr-MPRA study. In particular, the original study was enriched for regions which were putatively regulatory. If the method works well for other kinds of likely less functionally important regions, it would allow for a variety of interesting extensions.

For one thing, we can use our scores to see how enriched bases that are hypothesized to be higher in functional significance are for high importance. Disease-causing SNPs are one example of such a set of positions. More generally, there is recent work in weighted correction in situations where multiple hypotheses are tested. One can use incorporate the importance scores as prior side information in controlling false discovery rate, for example in GWAS studies.

Finally, there are several other datasets of possibly functionally-related annotations which can be compared against our model’s predictions. Several of these were explored in the Sharpr-MPRA study, including long terminal repeat elements (especially endogenous retroviral sequence 1 repeats) and endogenous chromatin state.

## 7 Conclusion

In this work we explore using convolutional neural networks to predict and understand transcriptional regulation. Using MPRA data published by Ernst et al. [3], we train our network to learn a mapping from 145-bp sequences to the output of the reporter assay for the sequence. After the model is trained, we generate per-base importance via *in silico* mutagenesis. To validate the model, we compare the generated importance scores to both transcription factor binding sites predicted by the CENTIPEDE algorithm [6] and evolutionary conserved regions [5].

Our model is able to both predict the regulatory score of a 145-bp sequence and identify functionally important nucleotides, offering a more powerful and flexible model than the PGM used in the Sharpr-MPRA study [3]. For example, the Sharpr computational model is explicitly tied to the experimental setup of the assay, because it models the overlap between 145-bp tiles; in contrast, our network learns from the tiling method used in the assay, but does not explicitly model it. In addition, the filters of the network are naturally interpreted as position weight matrices. Because the importance scores we produced successfully matched various other function-related annotations of genome positions, we believe it shows promise as a predictive tool in informing likely candidate locations for other predictive tasks.

